# Unaltered T cell responses to common antigens in individuals with Parkinson’s disease

**DOI:** 10.1101/2022.04.05.487195

**Authors:** Gregory P. Williams, Kaylin Muskat, April Frazier, Yaqian Xu, José Mateus, Alba Grifoni, Ricardo da Silva Antunes, Daniela Weiskopf, Amy W. Amara, David G. Standaert, Jennifer G. Goldman, Irene Litvan, Roy N. Alcalay, David Sulzer, Cecilia S. Lindestam Arlehamn, Alessandro Sette

## Abstract

**Background and Objectives:** Parkinson’s disease (PD) is associated with a heightened inflammatory state, including activated T cells. However, it is unclear whether these PD T cell responses are antigen specific or more indicative of generalized hyperresponsiveness. Our objective was to measure and compare antigen-specific T cell responses directed towards antigens derived from commonly encountered human pathogens/vaccines in patients with PD and age-matched healthy controls (HC).

**Methods:** Peripheral blood mononuclear cells (PBMCs) from 20 PD patients and 19 age-matched HCs were screened. Antigen specific T cell responses were measured by flow cytometry using a combination of the activation induced marker (AIM) assay and intracellular cytokine staining.

**Results:** Here we show that both PD patients and HCs show similar T cell activation levels to several antigens derived from commonly encountered human pathogens/vaccines in the general population. Similarly, we also observed no difference between HC and PD in the levels of CD4 and CD8 T cell derived cytokines produced in response to any of the common antigens tested. These antigens encompassed both viral (coronavirus, rhinovirus, respiratory syncytial virus, influenza, cytomegalovirus) and bacterial (pertussis, tetanus) targets.

**Conclusions:** These results suggest the T cell dysfunction observed in PD may not extend itself to abnormal responses to commonly encountered or vaccine-target antigens. Our study supports the notion that the targets of inflammatory T cell responses in PD may be more directed towards autoantigens like α-synuclein (α-syn) rather than common foreign antigens.

## Introduction

Parkinson’s disease (PD), the second most common neurodegenerative disease behind Alzheimer’s disease (AD), affects over 6 million people globally. PD symptomology is typically characterized by motor dysfunction including bradykinesia, tremor, and rigidity. However, non-motor symptoms often present long before clinical diagnosis and include loss of smell, disturbed sleep (e.g., REM sleep behavior disorder), constipation, and depression^1^. The major pathologies believed to underlie these PD symptoms are the degeneration of neurons (especially dopaminergic neurons in the substantia nigra) as well as the excessive accumulation and aggregation of the neuronal protein α-synuclein (α-syn)^2^.

Alongside the hallmark neuronal pathology observed in PD is the observation of increased pro-inflammatory markers among several different cell types and spanning multiple tissues^3^. This includes both microgliosis and astrogliosis in the central nervous system (CNS)^4,5^, as well as dysregulated monocyte populations in circulating blood^6,7^. However, the heightened PD inflammatory state is not limited to innate immune cells; increased CD4 and CD8 T cell infiltration in PD postmortem brain and abnormal blood T cell frequencies, compared to healthy controls (HC), have also been observed in some studies^8-10^. More recent findings have expanded on the potential inflammatory nature of T cells in PD and shown that some PD patients have active α-syn-specific T cell responses^11,12^. A follow-up study showed that T cell responses in PD patients were associated with early PD and a longitudinal study of a single case showed that α-syn-specific T cell responses were present years before the individual displayed symptoms^13^. Furthermore, the α-syn evoked T cell responses reached their highest magnitude around the same time as that individual acquired a varicella-zoster virus (shingles) infection. Intriguingly, it has been observed that individuals on anti-TNF treatment are at a lower risk of developing PD^14^. It is, however, unclear whether these results are reflective of a general heightened inflammatory state and altered global immune function in PD, or more explicitly reflective of T cell responses specific to α-syn and potentially other self-antigens.

We set out to test whether there is a general increased T cell responsiveness in individuals with PD compared to age-matched HC. We utilized different cell-based assays to determine the magnitude, specificity, and type of T cell response to commonly encountered pathogen and vaccine antigens. These common antigens included common cold coronaviruses, rhinovirus, respiratory syncytial virus, influenza, cytomegalovirus, varicella-zoster virus, pertussis, and tetanus toxoid. Using the activation induced marker (AIM) assay^15-17^ and intracellular cytokine staining (ICS) as a readout for T cell activation, we exposed both PD patient and age-matched HC peripheral blood mononuclear cells (PBMCs) to pathogen-specific peptide pools that have been previously shown to be targets of T cell responses. We found no difference in antigen-specific T cell activation between HC and individuals with PD. These results suggest that the inflammatory profile associated with PD does not influence the immune response to commonly encountered pathogens and vaccine targets.

## Methods

### Ethics statement

All participants provided written informed consent for participation in the study. Ethical approval was obtained from the Institutional Review Boards at La Jolla Institute for Immunology (LJI; Protocol Nos: VD-124 and VD-118), Columbia University Irving Medical Center (CUMC; protocol number IRB-AAAQ9714 and AAAS1669), University of California San Diego (UCSD; protocol number 161224), Rush University Medical Center (RUMC; Office of Research Affairs No.16042107-IRB01), Shirley Ryan AbilityLab Northwestern University (protocol number STU00209668-MOD0005), and the University of Alabama at Birmingham (UAB; protocol number IRB-300001297).

### Study subjects

39 participants were recruited for this study, 20 individuals diagnosed with PD and 19 age-matched healthy subjects. These subjects were recruited across multiple sites which consisted of Columbia University Irving Medical Center (CUMC) (HC n=1 and PD n=6), La Jolla Institute for Immunology (LJI) (HC n=11 and PD n=10), University of California San Diego (HC n=3 and PD=2), Rush University Medical Center (RUMC) (HC n=1), Shirley Ryan AbilityLab Northwestern University (HC n=1 and PD n=2), and the University of Alabama at Birmingham (UAB) (HC n=2). The characteristics of the enrolled subjects are detailed in Table 1.Subjects with idiopathic PD and HCs were recruited by the Movement Disorders Clinic at the Department of Neurology at CUMC, by the clinical core at LJI, by the Parkinson and Other Movement Disorder Center at UCSD, by movement disorder specialists at the Parkinson’s disease and Movement Disorders program at Shirley Ryan AbilityLab, and by movement disorder specialists at UAB Movement Disorders Clinic. Inclusion criteria for PD patients consisted of i) clinically diagnosed PD with the presence of bradykinesia and either resting tremor or rigidity ii) PD diagnosis between ages 35-80 iii) history establishing dopaminergic medication benefit, and iv) ability to provide informed consent. Exclusion criteria for PD were atypical parkinsonism or other neurological disorders, history of cancer within past 3 years, autoimmune disease, and chronic immune modulatory therapy. Age and sex matched HC were selected on the basis of i) age 45-85 and ii) ability to provide informed consent. Exclusion criteria for HC was the same as PD except for the addition of self-reported PD genetic risk factors (i.e., PD in first-degree blood relative). For the LJI cohort, PD was self-reported. Individuals with PD recruited at CUIMC, UCSD, and Northwestern all met the UK Parkinson’s Disease Society Brain Bank criteria for PD.

**Table 1:** Cohort General Information

|  | HC | PD |  |
| --- | --- | --- | --- |
|  | n = 19 | n = 20 | p value |
| **Sex (male/female)** |  |  |  |
| Male | 9 | 10 | N/A |
| Female | 10 | 10 | N/A |
| **Age (mean, SD):** |  |  |  |
| Total | 65.00 ± 5.98 | 65.40 ± 5.96 | 0.83 |
| Male | 64.11 ± 5.15 | 65.10 ± 5.78 | 0.7 |
| Female | 65.80 ± 6.81 | 65.70 ± 6.47 | 0.97 |
| **Caucasian (%):** | 89 (17/19) | 85 (17/20) |  |
| **Time Since PD Diagnosis (mean, range):** |  |  |  |
| Total | N/A | 8.05 (1-28) |  |
| Male | N/A | 7.90 (1-18) |  |
| Female | N/A | 8.20 (2-28) |  |
| **UPDRS Part III<sup>a</sup> (mean, SD):** |  |  |  |
| Total | N/A | 18.5 ± 5.73 |  |
| Male | N/A | 17 ± 6.96 |  |
| Female | N/A | 21 ± 1.73 |  |
| **MoCA Score (mean, SD):** |  |  |  |
| Total | N/A | 25.36 ± 2.46 |  |
| Male | N/A | 24.08 ± 2.06 |  |
| Female | N/A | 25.36 ± 3.27 |  |

<sup>a</sup>UPDRS Part III collected at CUMC, UAB, RUMC, and UCSD.
<sup>b</sup>MoCA collected for PD at CUMC, LJI, and UCSD.

### PBMC isolation

Venous blood was collected from each participant in either heparin or EDTA containing blood bags/tubes. PBMCs were isolated from whole blood by density gradient centrifugation using Ficoll-Paque plus (GE #17144003). Blood was first spun at 1850 rpm for 15 mins with brakes off to remove plasma. Plasma depleted blood was then diluted with RPMI and 35 mL of blood was carefully layered on tubes containing 15 mL Ficoll-Paque plus. These tubes were then centrifuged at 1850 rpm for 25 mins with brakes off. The interphase cell layer resulting from this spin were collected, washed with RPMI, counted, and cryopreserved in 90% v/v FBS and 10% v/v DMSO and stored in liquid nitrogen until tested. The detailed protocol for PBMC isolation can be found at protocols.io (https://dx.doi.org/10.17504/protocols.io.bw2ipgce).

### Antigen pools

Our lab routinely identifies CD4 and CD8 T cell epitopes, as well as uses existing data in the Immune Epitope Database and Analysis Resource (IEDB^18^) to develop peptide “megapools”. The utilization of “megapools” allows the ability to test a large number of epitopes spanning multiple HLA-types allowing a breadth of responses in immuno-diverse populations. Peptides were synthesized commercially as crude material by TC Peptide Lab (San Diego, CA). Lyophilized peptide products were dissolved in dimethyl sulfoxide (DMSO) at a concentration of 20 mg/mL and their quality was spot checked by mass spectrometry. Overlapping and predicted class II peptides were combined to form antigen “pools” for all of the respective antigens tested. Composition of the peptide pools used in this study have been previously defined, characterized, and are listed in Table 2. Note: Flu-Other references influenza antigens derived from flu proteins “other” than HA.

**Table 2:**
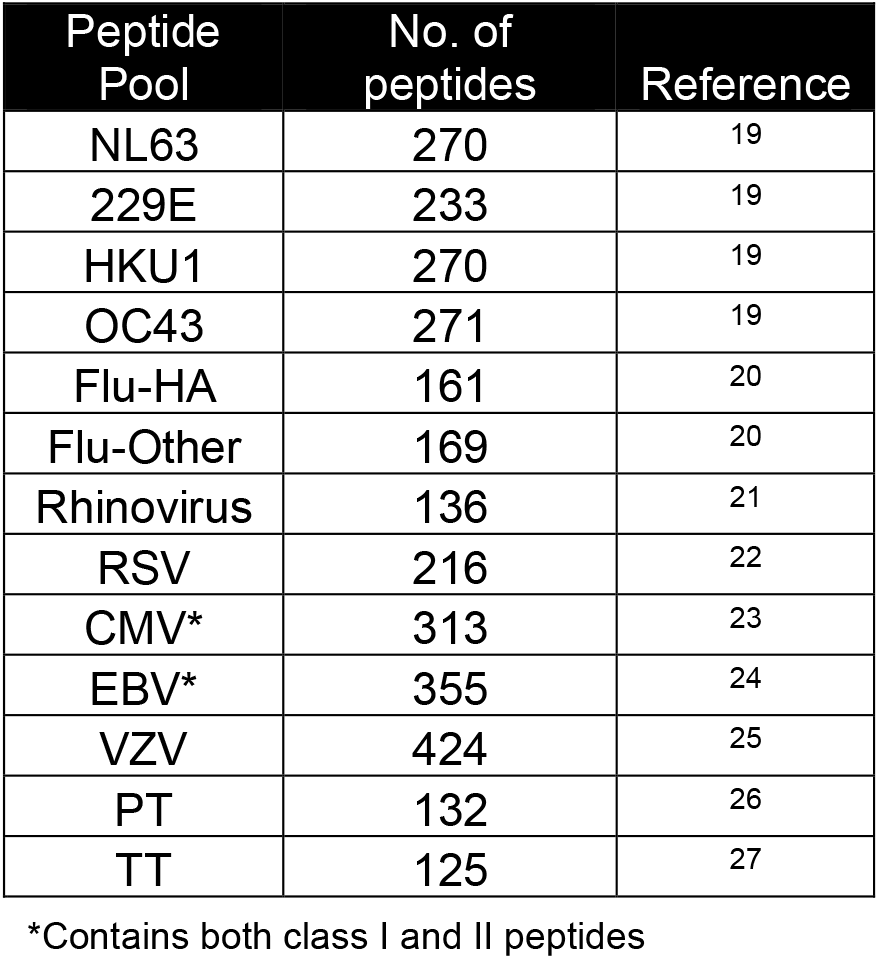
Antigen pools compositions

| Peptide Pool | No. of peptides | Reference |
| --- | --- | --- |
| NL63 | 270 | 19 |
| 229E | 233 | 19 |
| HKU1 | 270 | 19 |
| OC43 | 271 | 19 |
| Flu-HA | 161 | 20 |
| Flu-Other | 169 | 20 |
| Rhinovirus | 136 | 21 |
| RSV | 216 | 22 |
| CMV* | 313 | 23 |
| EBV* | 355 | 24 |
| VZV | 424 | 25 |
| PT | 132 | 26 |
| TT | 125 | 27 |

\*Contains both class I and II peptides

### Activation-induced marker (AIM) assay

PBMCs were thawed, washed, and counted for viability using the trypan blue exclusion method. A total of 1×10^6^ cells per subject/condition were plated and cultured (after a 15min pre-incubation with anti-CD40 (Miltenyi, clone HB14, RRID:AB_2660897) in the presence of each of the previously described peptide pools (at a concentration of 1μg/mL), PHA (10 μg/mL), or medium containing an equivalent amount of DMSO as what is present in the pool stimulation in a 96 well U-bottom plate for 24 hrs at 37 °C, 5% CO2. After incubation, cells were then washed and stained with a cocktail of antibodies that included CXCR5-BUV395 (BD, clone RF8B2, RRID:AB_2740008), CD8-BUV496 (BD, clone RPA-T8, RRID:AB_2870223), CD3-BUV805 (BD, clone UCHT1, RRID:AB_2739277), CD45RA-BV421 (Biolegend, clone HI100, RRID:AB_10965547), Fixable Viability Dye eFluor506 (eBioscience), CD14-V500 (BD, clone M5E2, RRID:AB_10611856), CD19-V500 (BD, clone HIB19, RRID:AB_105623910), CD4-BV605 (BD, RPA-T4, RRID:AB_2744420), CD38-BV786 (BD, clone HIT2, RRID:AB_2738515), CCR7-FITC (Biolegend, clone G043H7, RRID:AB_10916386), CD40L-PerCP-ef710(eBioscience, clone 24-31, RRID:AB_10670357), CD69-PE (BD, clone FN50, RRID:AB_395916), PD-1-PE-Dazzle594 (Biolegend, clone EH12.2H7, RRID:AB_2563659), OX40-PE-Cy7 (Biolegend, clone Ber-ACT35, RRID:AB_10901161), CD137-APC (Biolegend, clone 4B4-1, RRID:AB_830672), and HLA-DR-AF700 (eBioscience, clone LN3, RRID:AB_1907427) for 30 min at 4 °C. Afterwards, cells were washed and resuspended in 100 μL PBS for acquisition.

Cells were acquired on a Bio-Rad ZE5 cell analyzer, and further analysis was done using FlowJo software. Per previous publications^15,16^, quantification of live, single antigen-specific CD4^+^ and CD8^+^ T cells was determined as a percentage of their OX40/CD137 and CD69/CD137 double expression (AIM^+^), respectively. Antigen specific AIM^+^ CD4 and CD8 T cell signals were background subtracted by their corresponding DMSO negative controls, with the minimal DMSO level set to 0.005%. The limit of detection (LOD) for the AIM assay was calculated by multiplying the upper confidence interval of the geometric mean of all AIM^+^ DMSO samples by 2. The detailed protocol for the AIM staining assay can be found at protocols.io (submitted, pending approval).

### Intracellular cytokine assa

For the intracellular cytokine assay, PBMCs were processed and stimulated in the same way as the AIM assay (described above), but with the following modifications: After the 24hr incubation cells were then given a combination of Golgi-Stop (BD) and Golgi-Plug (BD) for 4 additional hours of culture. Afterwards the cells were surface and viability stained with CD3-BUV395 (BD, clone UCHT1, RRID:AB_2744387), CD8-BUV661 (BD, clone RPA-T8, RRID:AB_2871032), CD16-BV510 (Biolegend, clone 3G8, RRID:AB_2562085), CD14-BV510 (Biolegend, clone 63D3, RRID:AB_2716229), CD20-BV510 (Biolegend, clone 2H7, RRID:AB_2561941), CD45RA-BV750 (Biolegend, clone HI100, RRID:AB_2563813), CD4-BV711 (BD, clone RPA-T4, RRID:AB_2740432), CCR7-Pe-Cy7 (Biolegend, clone G0343H7, RRID:AB_11126145), and Fixable LIVE/DEAD Blue (Thermo Fisher) and incubated for 30 min at 4 °C. Cells were then washed, fixed (4% paraformaldehyde/PBS buffer), and then subsequently stained for intracellular cytokines with IL-4-BUV737 (BD, clone MP4-25D2, RRID:AB_2870157), IL-17-BV785 (Biolegend, clone BL168, RRID:AB_2566765), TNF-eFluor450 (eBioscience, clone Mab11, RRID:AB_2043889), IFNγ-FITC (eBioscience, clone 4S.B3, RRID:AB_465415), IL-2-BB700 (BD, clone MQ1-17H12, RRID:AB_2744488), IL-10-PE-Dazzle594 (Biolegend, clone JES3-19F1, RRID:AB_2632783), and CD40L-APC-eFluor780 (LifeTech, clone 24-31, RRID:AB_1603203) with saponin containing buffer at room temperature for 30 mins. Afterwards, cells were washed and resuspended in 100 μL PBS for acquisition.

Cells were acquired on a Bio-Rad ZE5 cell analyzer, and further analysis was done using FlowJo software. Similar to AIM analysis, quantification of live, singlet, cytokine producing CD4 cells were determined as a percentage of their CD40L^+^ Cytokine^+^ expression. CD40L/Cytokine double positive signals were subtracted by their corresponding DMSO negative controls. The limit of detection (LOD) for the ICS assay was calculated by multiplying the upper confidence interval of the geometric mean of all Cytokine^+^ DMSO samples by 2. The detailed protocol for the intracellular cytokine staining assay can be found at protocols.io (submitted, pending approval).

### Data analysis

Lymphocytes were analyzed using a standardized gating method (see Supplemental Fig. 1 for detailed gating strategy) used within the lab. On top of the previously described signal background subtraction in both AIM and ICS assays, another quality-control measure employed in this study was the exclusion of samples that did not have a ≥20% AIM/Cytokine^+^ signal to the PHA-positive control stimuli. Statistical analyses were performed and graphs created using GraphPad Prism’s descriptive statistics and two-tailed Mann Whitney tests.

## Results

### T cell reactivity to common antigens in HC and PD PBMCs

We utilized PBMCs from a cohort of age and sex-matched 19 HC and 20 PD individuals recruited from multiple sites (see Table 1 for detailed subject characteristics).

PBMCs were stimulated with antigenic peptide pools specific for coronaviruses NL63, 229E, HKU1, OC43, influenza virus (FLU-HA, FLU-Other), rhinovirus, respiratory syncytial virus (RSV), cytomegalovirus (CMV), varicella-zoster virus (VZV), *Bordetella pertussis* (PT), and tetanus toxoid (TT) antigens —see Table 2 for more detailed information on peptide pool compositions. Antigen pools tested were grouped into general classifications to aid in data viewing (e.g., Common Coronavirus, Lower Respiratory Tract Viruses). For CMV and EBV pools we also measured CD8-specific reactivity in addition to CD4 T cell responses (as these pools contain both class I and class II peptides). The antigen-specific T cell responses were measured using a flow cytometry activation-induced marker (AIM) assay^15-17^ and antigen-specificity was determined as percentage of CD4^+^ cells expressing both OX40^+^ and CD137^+15,16^, or CD8^+^ cells expressing both CD69^+^ and CD137^+15,16^ (Fig. 1a-d and Supplemental Fig. 1 for AIM gating strategy). No significant difference in the percentage of antigen-specific CD4 T cells between HC and PD was observed for any of the antigens tested. Similarly, we observed no difference in the CMV or EBV-specific CD8 AIM response between HC and PD (Fig 1e). Additionally, we also measured the upregulation of other combinations of activation-associated markers and found no differences in responses between HC and PD (Supplemental Fig. 2).

**Fig. 1:**
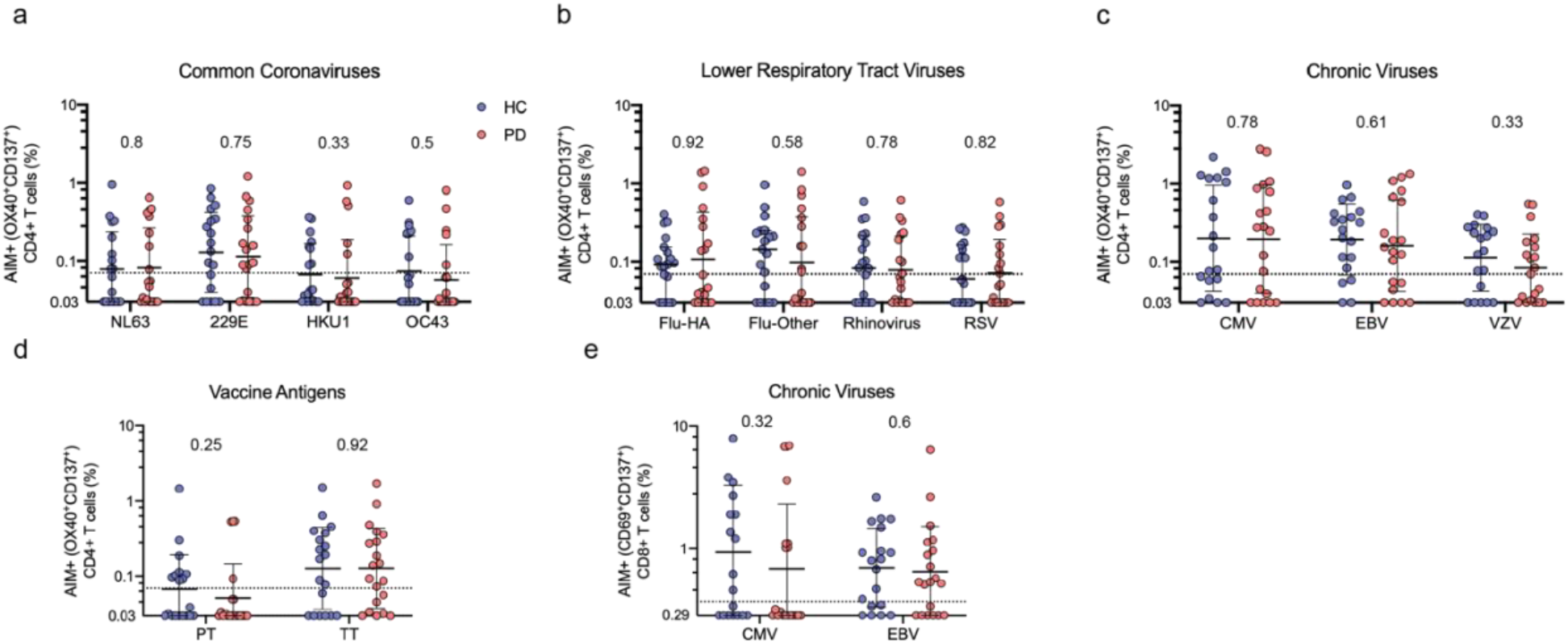
Antigen specific T cell responses to common antigens in HC and PD patient’s PBMCs. a-d) AIM responses (OX40^+^ CD137^+^) in CD4 T cells and e) CD8 T cells (CD69^+^ CD137^+^) after stimulation to previously defined antigenic peptide pools. PBMCs from HC (blue circles) and PD (red circles) were stimulated with 1 ug/mL of NL63, 228E, HKU1, OC43, Flu-HA, Flu-Other, Rhinovirus, RSV, CMV, EBV, VZV, PT, or TT for 24 hrs, stained, and run on a Bio-Rad ZE5 flow cytometer. e) AIM responses (CD69^+^ CD137^+^) in CD8 T cells. Each dot represents an individual subject, HC n=19, PD n=20. Two-tailed Mann-Whitney test; geometric mean with standard deviation is displayed. Dashed line indicates threshold of positivity (TP).

### T cell derived cytokine response to common antigens in HC and PD PBMCs

As described above, we did not observe any significant differences between antigen-specific T cell activation in HC and PD individuals. To expand this observation, we next utilized a different assay system and addressed whether there is a difference in the specific cytokines produced between the two cohorts. PBMCs were stimulated with the same antigenic peptide pools used in the previous experiment and then the specific cytokine production was measured using an intracellular cytokine staining assay where we measured IFNγ, TNFα, IL-2, IL-17a, IL-4, and IL-10 to represent the majority of Th subsets. In terms of Th1 associated cytokine responses, there was no statistical difference between HC and PD CD4^+^, CD40L^+^, IFNγ producing T cells after stimulation with any of the antigen pools tested (Fig. 2a). Some marginal reductions were seen for certain antigen/cytokine combinations, such as a lower CD4+ TNFα-production in PD subjects in response to the 229E coronavirus and VZV antigen pool (Fig. 2b), and reduced IL-2 production against the same common cold corona virus 229E antigen pool (Fig. 2c). These decreases were however not significant after considering corrections for multiple comparisons (Bonferroni corrected critical value of 0.003). No detection or low levels of IL-4, IL-10, and IL-17a, along with CD8+ IFNγ or TNFα were noted, with no difference in levels between HC and PD individuals (data not shown). Overall, we observed a generalized trend of lower cytokine responses in PD individuals and more specifically to the 229E common cold coronavirus strain, however these differences would not meet significance if corrected for multiple comparisons. Taken together, these results indicate that there is no difference between HC and PD in regard to antigen-specific T cell reactivity nor the types of cytokines produced to several commonly encountered antigens.

**Fig. 2:**
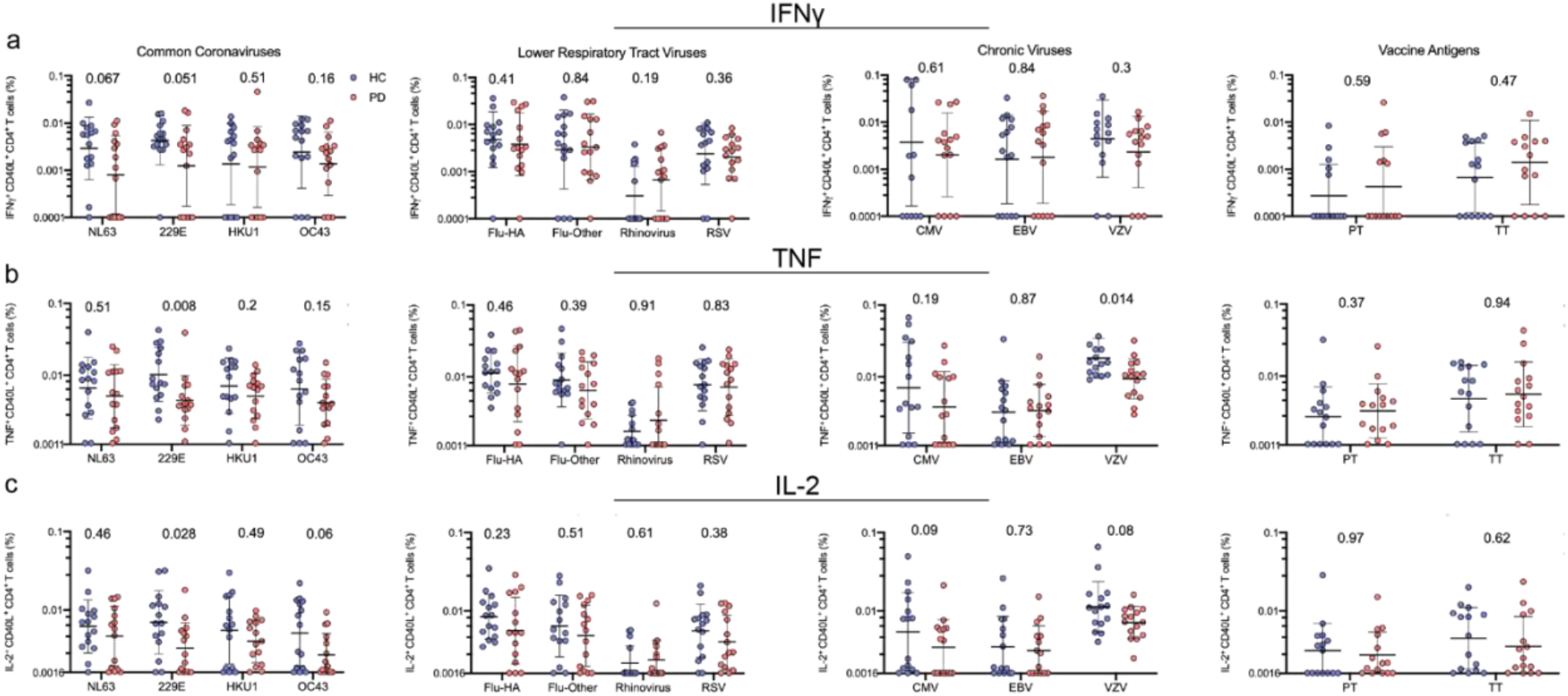
T cell derived cytokine responses to common antigens in HC and PD patients PBMCs. a-d) Intracellular cytokine responses in CD4 T cells after stimulation to previously defined antigenic peptide pools. PBMCs from HC (blue circles) and PD (red circles) were stimulated with 1 ug/mL of NL63, 228E, HKU1, OC43, Flu-HA, Flu-Other, Rhinovirus, RSV, CMV, EBV, VZV, PT, or TT for 24 hrs, stained, and run on a Bio-Rad ZE5 flow cytometer. Cells were gated on live, singlet CD4^+^ populations and then on different cytokine populations (a: IFNγ/CD40L^+^ b: TNFα/CD40L^+^, c: IL-2/CD40L^+^). Each dot represents an individual subject, PD n=20, HC n=19. Two-tailed Mann-Whitney test; geometric mean with standard deviation is displayed.

## Discussion

This study investigated whether individuals with PD have T cell responses that are different compared to age-matched healthy controls after stimulation with commonly encountered antigens. The common antigens tested encompassed multiple types of pathogens including common cold coronavirus, influenza, and herpesvirus. Both T cell activation and corresponding cytokine output were considered and we found no significant differences between the two cohorts. Our findings suggest that the hyper-inflammatory state reported in PD does not influence antigen-specific T cell responses to common pathogens consistently encountered either through natural exposure or vaccination. This observation is in agreement with the notion that the targets of peripheral T cell inflammation observed in PD are more directed towards autoantigens such as α-synuclein^11-13,28^.

Multiple groups have observed and reported on dysregulated and pro-inflammatory T cell responses detected in individuals with PD^9,10^,29,30. These findings include observations of increased antigen experienced T cells^29^ as well as increases in T cell derived IFN-γ,^10^ IL-4^10^, and IL-17^9,10^. However, these reports measured baseline T cell frequencies or cytokine responses following polyclonal stimulation such as PMA/Ionomycin. While these methods are informative in establishing the steady state of PD T cell composition and activation potential, they fall short of uncovering what targets are seemingly driving the pro-inflammatory T cell responses observed in the peripheral blood of PD patients. The work described here was aimed to determine if the baseline T cell hyper-responsiveness established by previous groups extends to a wide variety of antigens across several pathogens and vaccine antigens. For the pathogens we tested, antigen-specific T cell activation/cytokine production in peripheral blood was not different in the PD and HC cohorts.

A potential link between infectious disease and development of Parkinson’s disease has long been reported on across multiple different pathogens. An outbreak of encephalitic lethargica was reported around the time of the 1918 flu pandemic and transient neurological sequelae driven PD symptoms were associated with early stages of influenza infection^31^. Links between influenza infection and subsequent risk of developing PD have been debated throughout the years^32-35^. Most recently, an observational study found an increase in PD incidence following clinical influenza infection^36^. Two potential mechanisms that could link infection and development of PD would be the direct insult of the infection to the nigrostriatal system or the infection leading to an over-activation of the immune system (a common feature of PD). However, the data presented here does not suggest an increased influenza-specific T cell response in PD patients compared to healthy controls. Another pathogen tested in this study, Epstein-Barr virus has previously been linked to PD^37-39^ and EBV has been heavily implicated in development of another CNS disease, multiple sclerosis^40^. Lastly, it has been hypothesized that coronavirus infection may be involved in PD pathogenesis as increased CSF coronavirus antibodies have been previously been observed in PD patients^41^ along with the fact that a coronavirus strain in mice was observed to infect and deteriorate their basal ganglia^42^. To our knowledge, this is the first study of its kind to measure antigen-specific T cell responses in patients with PD and HC to pathogens previously linked to PD (influenza, coronavirus, and EBV), as well as previously unassociated infectious species and vaccine antigens (rhinovirus, pertussis, and tetanus).

In conclusion, though there are multiple reports detailing a baseline hyper-inflammatory state in PD, our results shown here suggest that this hyper-responsiveness does not apply to antigen-specific T cell responses to many commonly encountered antigens. Given the fact that there have been multiple associations with PD and infectious triggers, it may be that other facets of the immune system (auto-antibodies, bystander damage due to general inflammation) could be of more importance to these epidemiological connections than antigen-specific T cell responses. The only increased antigen-specific T cell response observed thus far in PD has been to α-syn, a neuro-antigen. More work is needed in exploring if other relevant neuro-antigens might be the target of PD T cells and could better explain the sources of increased inflammation present in Parkinson’s disease.

## Acknowledgement

The authors want to sincerely thank all of the generous individuals who donated samples for the study. We are also grateful to the La Jolla Institute for Immunology’s clinical core for processing the blood samples.

## Author contributions

KM performed the experiments, GPW, KM analyzed the data, AF, YX recruited participants and AWA, DGS, JGG, IL, RNA performed clinical evaluations. JM, AG, RA, and DW designed, characterized, and provided epitope pools. GPW wrote the manuscript, DS, CLA, ALS des{igned and discussed the data. All authors read, edited, and approved the manuscript before submission.

