## Supplemental Figures 1,2 for "Unaltered T cell responses to common antigens in individuals with Parkinson’s disease"

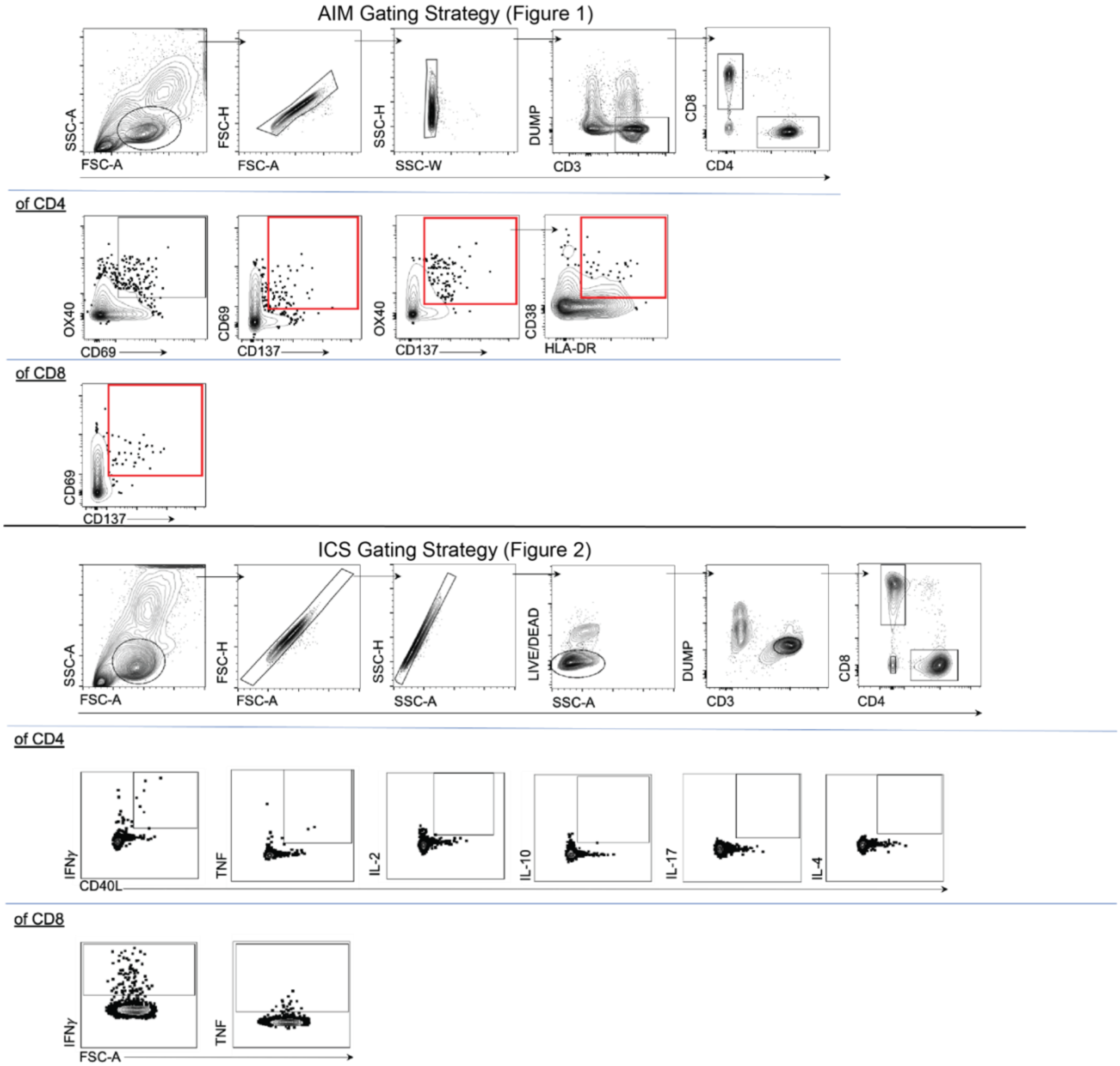


**Supplemental Fig. 1 Gating strategies for AIM and ICS analyses:** Top panel, representative strategy used to define AIM+ cells described in Figure 1. Briefly, post-incubated PBMCs were stained and then gated for lymphocytes, singlets, live, CD3+, and then CD4+ or CD8+. For CD4, double positive expression of either OX40, CD69, and CD137 were used to define AIM+ cells (with OX40 and CD137 being the main population). For CD8, double positive expression of CD69/CD137 was used to define AIM+ cells. Lower panel, similar to the AIM gating strategy, for ICS (related to Figure 2), lymphocyte, singlets, live, CD3+, and CD4+ or CD8+ T cells were selected. For CD4 T cells, double positive expression for cytokine (IFN$\gamma$, TNF, etc.) and CD40L was used to determine antigen-specific cytokine producing T cells. For CD8, just single expression of cytokine was used.

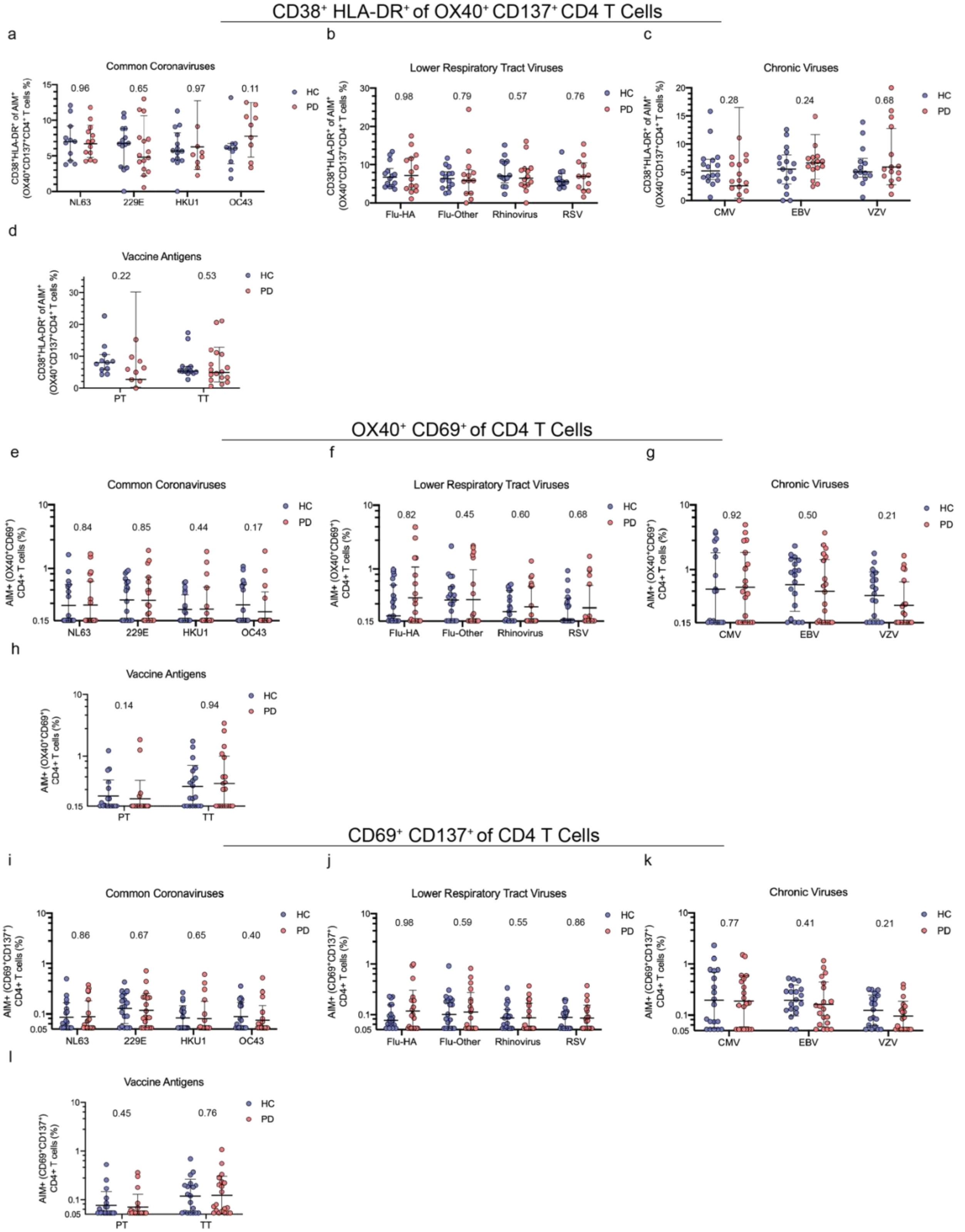
CD38+ HLA-DR+ of OX40+ CD137+ CD4 T Cells
a
Common Coronaviruses
0.96
0.65
0.97
0.11
NL63
229E
HKU1
OC43
HC
PD
b
Lower Respiratory Tract Viruses
0.98
0.79
0.57
0.76
Flu-HA
Flu-Other
Rhinovirus
RSV
c
Chronic Viruses
0.28
0.24
0.68
CMV
EBV
VZV
d
Vaccine Antigens
0.22
0.53
PT
TT
OX40+ CD69+ of CD4 T Cells
e
Common Coronaviruses
0.84
0.85
0.44
0.17
f
Lower Respiratory Tract Viruses
0.82
0.45
0.60
0.68
g
Chronic Viruses
0.92
0.50
0.21
h
Vaccine Antigens
0.14
0.94
CD69+ CD137+ of CD4 T Cells
i
Common Coronaviruses
0.86
0.67
0.65
0.40
j
Lower Respiratory Tract Viruses
0.98
0.59
0.55
0.86
k
Chronic Viruses
0.77
0.41
0.21
l
Vaccine Antigens
0.45
0.76

**Supplemental Fig. 2 Additional AIM population levels between HC and PD to common antigens**: PBMCs from HC (blue circles) and PD (red circles) were stimulated with 1 ug/mL of NL63, 228E, HKU1, OC43, Flu-HA, Flu-Other, Rhinovirus, RSV, CMV, EBV, VZV, PT, or TT for 24 hrs, stained, and run on a Bio-Rad ZE5 flow cytometer. a-d) Quantification of recently activated responses ($CD38^+$ HLA-$DR^+$% of $OX40^+$ $CD137^+$ cells in Fig. 1) to the common antigens tested. Additional populations are displayed in e-h) $OX40^+$ $CD69^+$ $AIM^+$ CD4 T cell quantification and i-l) $CD69^+$ $CD137^+$ $AIM^+$ CD4 T cell quantification. Each dot represents an individual subject, PD n=20, HC n=19. Two-tailed Mann-Whitney test; geometric mean with standard deviation is displayed.
